# Dopamine response gene pathways in dorsal striatum MSNs from a gene expression viewpoint: cAMP-mediated gene networks

**DOI:** 10.1101/757500

**Authors:** Vladimir Babenko, Anna Galyamina, Igor Rogozin, Dmitry Smagin, Natalia Kudryavtseva

## Abstract

A mouse model of chronic social conflicts was used to analyze dorsal striatum neurons implicated in cAMP-mediated phosphorylation activation pathways specific for Medium Spiny Neurons (MSNs). Based on expression correlation analysis, we succeeded in dissecting Drd1- and Drd2-dopaminoceptive neurons (D1 and D2, correspondingly) gene pathways. We also found that D1 neurons feature previously reported two states, passive and active ones, represented in our analysis by distinct, negatively correlated gene clusters.

The correlation based gene pathways strongly corroborate the phosphorylation cascades highlighted in the previous studies, implying that the expression-based viewpoint corresponds to phosphorylation/dephosphorylation interplay in each type of neurons. Notably, D2 neurons showed the largest *Ppp1r1b* (encoding DARPP-32) expression modulation impact, implying that *Ppp1r1b* expression dynamics is mostly associated with neuroendocrine response mediated by *Penk/Pdyn* genes expression in D2 neurons.

We observed that under defeat stress in chronic social conflicts mice exhibited reduced motor activity as well as overall depression of dopamine-mediated MSNs activity, while aggressive mice exhibited motor hyperactivity and an increase in both D1-active phase and D2 MSNs genes expression.

Based on alternative transcript isoforms expression analysis, it was assumed that many genes (*Drd1, Adora1, Pde10, Ppp1r1b, Gnal*), specifically those in D1 neurons, apparently remain transcriptionally repressed via the reversible mechanism of promoter CpG island silencing, resulting in alternative promoter usage following profound reduction in their expression rate.

**Significance statement:** Medium Spiny Neurons (MSNs) comprise the main body of dorsal striatum neurons and represent dopaminoceptive GABAergic neurons. The cAMP- mediated cascade of excitation and inhibition responses involved in dopaminergic neurotransmission is crucial for neuroscience research due to its involvement in the motor and behavioral functions. In particular, all types of addictions are related to MSNs. Shedding the light on the mechanics of the above-mentioned cascade is of primary importance for this research domain. In this paper MSNs steady states will be elucidated based on pooled tissue RNA-Seq data not explicitly outlined before and connected with dynamic dopamine neurotransmission cycles.

## Introduction

The dorsal striatum is responsible for the regulation of motor activity and stereotypical behaviors and is also potentially involved in a variety of cognitive, reward and social hierarchy processes (Gerfen and Surmeier 2011; Lobo and Nestler 2011; Lee et al. 2018; Stagkourakis et al. 2018). Through afferent and efferent projections to associative, motor and sensorimotor cortical areas and other brain structures, dorsal striatum participates in the initiation and execution of movements, as well as in the regulation of muscle tone (Devan et al. 2011; Liljeholm and O’Doherty 2012).

Medium Spiny Neurons (MSNs) represent 95% of neuron population within the dorsal striatum in mice (Kemp and Powel 1971). Notably, MSNs are GABAergic neurons thatalso have dopamine/glutamate receptors in postsynaptic dendrites. The phosphorylation cascades throughout the dorsal striatum neurons play a central role in motion and emotion signals transduction (Surmeier et al. 2007). In particular, striatonigral “direct” and striatopallidal “indirect” pathways represent opposite excitatory and inhibitory signal transmissions, respectively, depending on MSN dopamine receptor types: D1-dopaminoceptive, or D2-dopaminoceptive (Gerfen et al. 1990; Ouimet et al. 1998).

Reciprocal protein phosphorylation/dephosphorylation cascades constitute a major regulatory mechanism of intracellular signal transduction. Its functioning in MSNs, where the corresponding gene pathways are highly expressed and coordinated, provides a vivid illustration. The networks comprise the pathways mediated by PKA and Cdk5 kinases for cAMP activated kinases, Mapk2-4 for mitogen-activated kinases, serine/threonine phosphatases PP1, PP2A, PP2B and two tyrosine phosphatases Ptpn5, Ptpn7 (Surmeier et al. 2007).

Due to the specifics of dephosphorylation machinery regulation, multiple inhibitor subunits are recruited in the phosphatase complex formation. In particular, protein phosphatase 1 (PP1) represented by three catalytic subunits alpha, beta, gamma, encoded by *Ppp1ca, Ppp1cb, Ppp1cc* genes, can associate with more than 100 inhibitor subunits (Eto 2009), while PP2A (2 catalytic subunits) inhibitors are represented by 15+distinct subunits, and PP2B (3 catalytic subunits) is regulated by four inhibitor subunit genes. However, it was found that only one inhibitor can bind the catalytic core at a time via the same catalytic site (Aggen et al. 2000). Thus, the repressor subunits play a pivotal role in regulating phosphatases in a tissue and stage specific manner.

Since protein phosphatases act in a wide range of cell types and are commonly associated with deactivation and ubiquitylation of proteins (Nguyen et al. 2013), *Ppp1r1b* encoding PP1 inhibitor subunit DARPP-32 was underlined as one of the rare neurospecific genes expressed at very high rates specifically in striatum medium spiny neurons and playing a crucial role in the ‘orchestration’ of neurotransmission (Svenningsson et al. 2004). *Ppp1r1b* is expressed in the upper bound of the expression range for dorsal striatum-related protein-encoding genes and is implicated as an ultimate factor of MSNs phosphorylation kinetics regulation (Scheggi et al. 2018). It has been proven to be involved in many pathophysiological processes (Albert et al. 2002; Svenningsson et al. 2005; Santini et al. 2007). Notably, this gene from the family of protein phosphatase inhibitors solely is directly associated with aggressive behavior as previous studies have revealed (Reuter et al. 2009).

In D1 MSNs cAMPs activate PKA which phosphorylates DARPP-32 on Thr34, transforming it into a of PP1 inhibitor (Stoof and Kebabian 1981; Nishi et al. 1997; Missale et al. 1998). Calcineurin (PP2B, its catalytic subunit expressed at the highest rates is encoded by *Ppp3ca*), is also expressed in D1 neurons, dephosphorylates DARPP-32 at Thr34, mediating DARPP-32 phosphorylated homeostasis state, in particular turning it off upon signal abrogation. Conversely, Cdk5, which is activated in D2-dopaminoceptive MSNs by Ca2+ provided by AMPA/NMDA receptors, phosphorylates DARPP-32 at Thr75, turning DARPP-32 into a PKA inhibitor (Nishi et al. 2002). The pathway is being regularly updated, and published elsewhere (Scheggi et al. 2018). Besides, MSNs express dorsal striatum specific Tyrosine phosphatase STEP (encoded by *Ptpn5*), which functions alternatively to serine/threonine phosphatases (Fitzpatrick and Lombroso 2011).

In the current study we investigated a mouse chronic social conflicts model with antagonizing emotional states and motor activity (hyperactivity and total immobility) (Smagin et al. 2018; Smagin et al.2019) to analyze the involvement of dorsal striatum neurons in cAMP-mediated phosphorylation activation pathways specific for Medium Spiny Neurons (MSNs). We aimed at assessing the overall profile of the D1/D2 neurons based on RNA-Seq expression profiling by considering the reported genes involved in phosphorylation kinetics in MSNs.

## Results

### Compilation of gene sets

To gain fuller insight into the D1/D2 MSNs we retrieved the annotated genes implicated in PP1-regulated posphorylation pathways in MSNs based on available literature (Nishi et al. 2002; Svenningsson et al. 2004; Nishi et al. 2011; Nishi et al. 2017; Yapo et al. 2017). The compiled core gene set comprising cAMP-mediated dopamine response genes expanded with NMDA glutamate receptor subunits *Grina*, *Grin1*-*Grin2* and MAP kinases is presented in Table 1. Their expression profiles are presented in Supplemental Table S1. We used *Cdk5r1*, neuron-specific activator of cyclin dependent kinase 5 (CDK5) p35 subunit, as a CDK5 activation marker. Three major serine-/threonine-specific phosphatases involved in MSN cascades are PP1, PP2A and PP2B. The PP1 catalytic core comprises 3 subunits (encoded by *Ppp1ca, Ppp1cb, Ppp1cc*), PP2A is represented by 2 catalytic subunits (*Ppp2ca* and *Ppp2cb*), and PP2B includes 3 subunits (*Ppp3ca*, *Ppp3cb, Ppp3cc*). For all three phosphatases the ‘alpha’ subunit exhibits the highest levels of expression (Supplemental Table S2), so these subunits were used as the major phosphatase gene markers, except for PP1 (all subunits were considered).

**Table 1.** Core genes set of cAMP-mediated dopamine response involved in the *Ppp1r1b* mediated phosphorylation cycles expanded with NMDA glutamate receptor set (*Grina, Grin1*-*Grin2*) and Map kinases genes.

| Gene symbol | Protein name | Description | STR-specific* |
| --- | --- | --- | --- |
| *Adcy5* | AC5 | adenylate cyclase 5 | yes |
| *Adora1* | A1A | adenosine A1 receptor | no |
| *Adora2a* | A2A | adenosine A2a receptor | yes |
| *Cdk5r1* | CDK5 | cyclin-dependent kinase 5, regulatory subunit 1 (p35) | no |
| *Drd1* | D1R | dopamine receptor D1 | yes |
| *Drd2* | D2R | dopamine receptor D2 | yes |
| *Gnai2* | Gi/o | guanine nucleotide binding protein (G protein), alpha inhibiting 2 | no |
| *Gnal* | Golf | guanine nucleotide binding protein, alpha stimulating, olfactory type | yes |
| *Gpr88* | STRG | G-protein coupled receptor 88 | yes |
| *Grin1* | NMDA1 | glutamate receptor, ionotropic, NMDA1 (zeta 1) | no |
| *Grin2a* | NMDA2A | glutamate receptor, ionotropic, NMDA2A (epsilon 1) | no |
| *Grin2b* | NMDA2B | glutamate receptor, ionotropic, NMDA2B (epsilon 2) | no |
| *Grin2c* | NMDA2C | glutamate receptor, ionotropic, NMDA2C (epsilon 3) | no |
| *Grin2d* | NMDA2D | glutamate receptor, ionotropic, NMDA2D (epsilon 4) | no |
| *Grina* | NMDARA1 | glutamate receptor, ionotropic, N-methyl D-aspartate-associated protein 1 (glutamate binding) | no |
| *Map2k1* | ERK1 | mitogen-activated protein kinase kinase 1 | no |
| *Map2k5* | ERK5 | mitogen-activated protein kinase kinase 5 | no |
| *Mapk1* | ERK2 | mitogen-activated protein kinase 1 | no |
| *Mapk3* | ERK1 | mitogen-activated protein kinase 3 | no |
| *Mapk7* | ERK5 | mitogen-activated protein kinase 7 | no |
| *Pde10a* | ADSD2 | phosphodiesterase 10A | yes |
| *Pdyn* | PPD | prodynorphin | no |
| *Penk* | PPE | preproenkephalin | yes |
| *Ppp1ca* | PP1 | protein phosphatase 1, catalytic subunit, alpha isoform | no |
| *Ppp1cb* |  | protein phosphatase 1, catalytic subunit, beta isoform | no |
| *Ppp1cc* |  | protein phosphatase 1, catalytic subunit, gamma isoform | no |
| *Ppp1r1b* | Darpp32 | protein phosphatase 1, regulatory (inhibitor) subunit 1B | yes |
| *Ppp2ca* | PP2A | protein phosphatase 2 (formerly 2A), catalytic subunit, alpha isoform | no |
| *Ppp3ca* | PP2B | protein phosphatase 3, catalytic subunit, alpha isoform (Calcineurin). | no |
| *Prkaca* | PKA | protein kinase, cAMP dependent, catalytic, alpha | no |
| *Ptpn5* | STEP | protein tyrosine phosphatase, non-receptor type 5 | yes |
| *Tac1* | PPTA | tachykinin 1 | yes |
| *Tac2* | PPTB | tachykinin 2 | no |

\*‘Striatum (STR) -specific’ column indicates genes maintaining striatum specific expression profile

### Basic confirmation of 4 clusters based on dorsal striatum data using the core genes set (Table 1)

We clustered the core dopamine response cAMP-mediated genes (Table 1) by Agglomerative Hierarchical Cluster (AHC) analysis (Fig. 1) and observed 4 distinct clusters. Strikingly, each of the clusters corresponds to a specific signal transduction cascade observed in cAMP-mediated response to dopamine, which can be ascribed to the key genes. In particular, there is a cluster comprising *Drd2* receptor distinct from one comprising *Drd1* receptor. Another cluster marks PKA-phosphorylation cascade *(Prkaca)*, and the fourth clusterrepresented by a single gene (*Tac2*) indicates the absence of dopamine signaling.

**Fig. 1.**
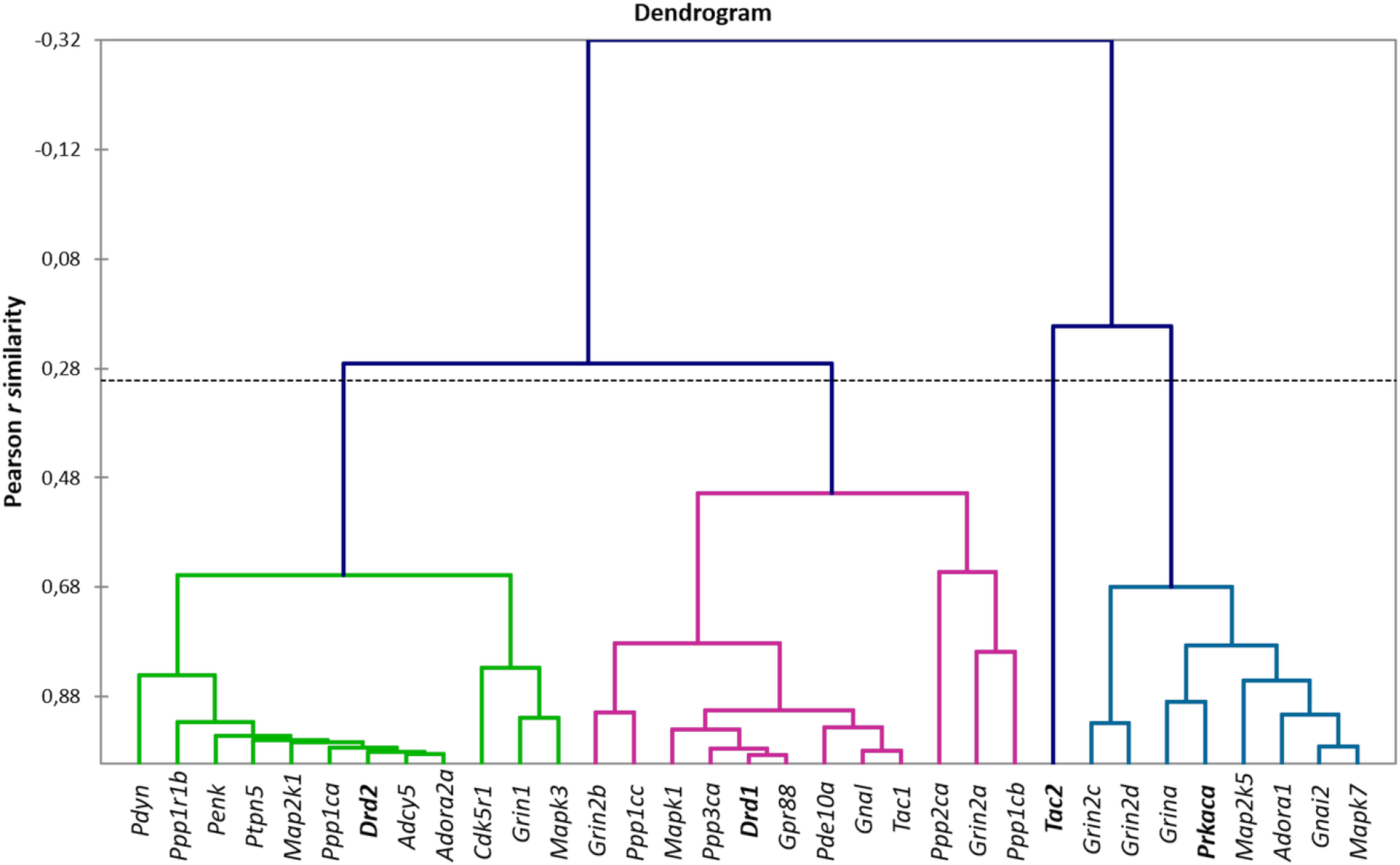
AHC analysis reveals 4 consequent clusters of a) D2-associated genes (green); b) D1-associated passive state genes (red); c) D1-associated active state genes (blue). Single gene corresponding to DA depletion is represented by *Tac2* expression.

**Figure 2.**
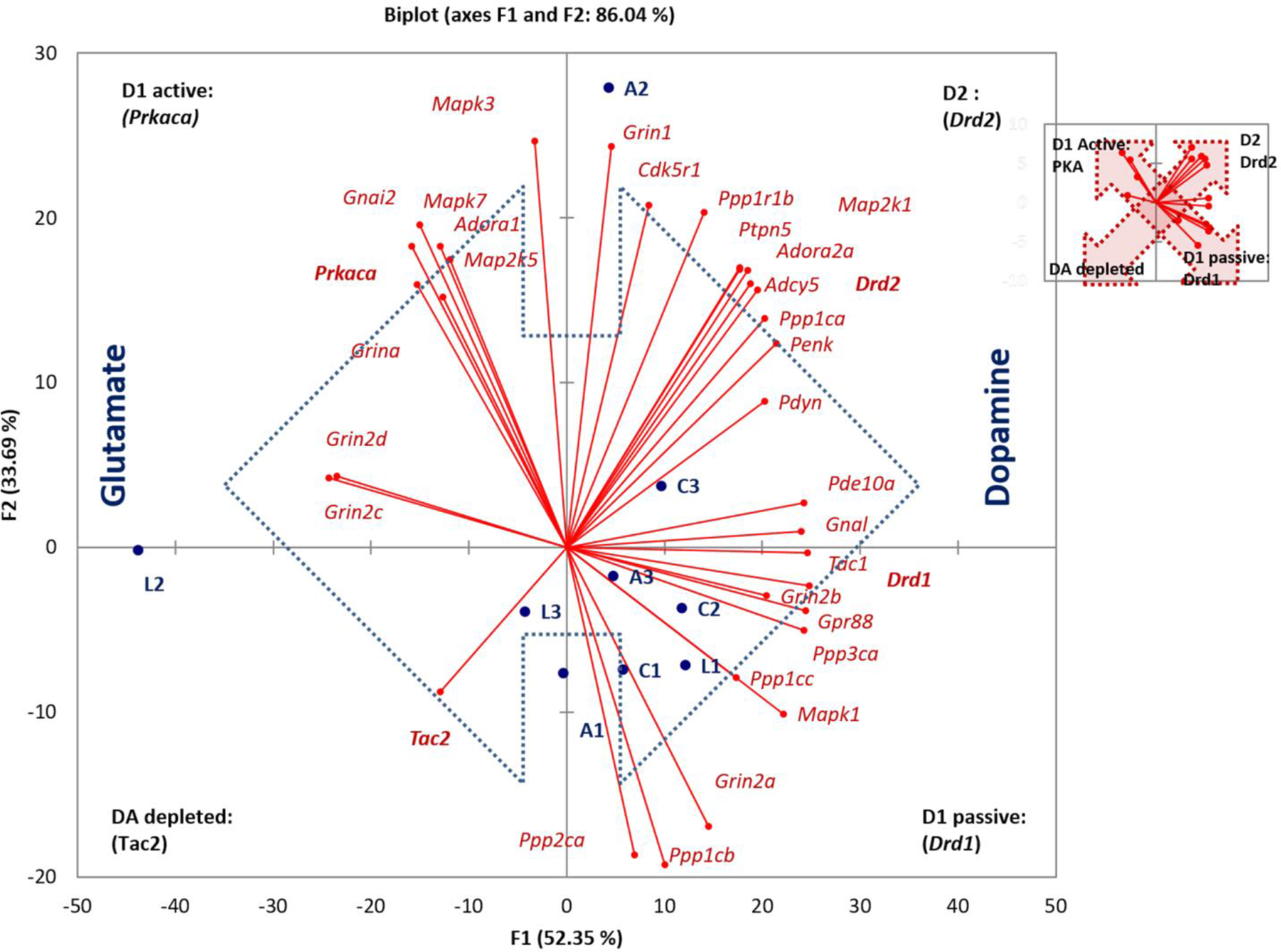
Major clusters detected by AHC and PCA: C1, C2, C3 – control; A1, A2, A3 – aggressive mice; L1, L2, L3 – losers, defeated mice. 4 quadrants are indicated with corresponding (D1/D2/DA depleted) states and marker genes. Inserted is a scheme of antagonistic clusters: the arrows depict alternative states of D1 MSN, and D2/DA-depleted states. Blue arrow indicates antagonistic gradient of dopamine and glutamate according to the corresponding receptors distribution. Inserted (small figure) is a scheme of antagonistic clusters: the arrows depict alternative states of D1 MSN, and D2/DA-depleted states. clusters of the D1 geneindicating its stable rather than passive state followed by a short firing time span upon activation represented by D1 active state as shown in (Yapo et al. 2017). The details of the PCA analysis are shown in Fig. 2. According to the genes in these clusters they were designated as D2 cluster, D1 passive, D1 active and DA depletion. Two major components are responsible for 86% of the variability in the expression of the genes evaluated, underscoring consistency and exhaustive representation of genes variation.

To gain an expanded view on the clusters elucidated by AHC we performed the Principal Components Analysis (PCA) on the same core genes set (Fig. 2). It has been confirmed that the gene clusters are highly synchronized (high correlation rate), each corresponding to a particular neuron type (D1/D2) according to their annotation in publications. Also, as follows from Fig. 2, we can observe antagonistic

#### Regulation of Ppp1r1b expression

It is known that in phosphorylated state DARPP-32 effectively inhibits both PP1 alpha (encoded by *Ppp1ca;* Hemmings et al. 1984) and PP1 gamma (encoded by *Ppp1cc;* Hemmings et al. 1989). As we can see from the plot in Fig.2 *Ppp1r1b* is located in the D2 cluster along with *Ppp1ca* encoding PP1 alpha subunit, which indicates that this cluster is a major expression site. However, another catalytic subunit of PP1, *Ppp1cc,* is located in D1 passive cluster (Fig. 2). Given *Ppp1cc* is committed specifically to D1 passive state (Fig. 2) one may expect its coexpression with *Ppp1r1b* in D1 neurons.

Similar to *Grin1* (major subunit of NMDA) committed to D2c luster, *Grin2a*-*d* subunits are distributed across distinct quadrants in Fig. 2: *Grin2a,b* are associated with D1 neurons passive state, while *Grina, Grin2d,c* are located in D1 neurons active state cluster.

### Expression rates of dorsal striatum-specific genes as compared to four other brain regions

Average expression rate of the genes has been analyzed in distinct clusters (described above) across 5 brain regions: hippocampus (HPC), hypothalamus (HPT), dorsal striatum (STR) ventral tegmental area (VTA), midbrain raphe nuclei (MRN). Several genes have been identified as the genes preferentially expressed in dorsal striatum. Expression profiles of the selected genes are shown in Fig.3a, b. They happen to be expressed specifically in D2 and D1 passive state clusters. As for D1 active cluster, we identified *Prkaca* (encoding one of the PKA subunits) and *Gnai2* as the genes with the lowest expression levels in dorsal striatum across 5 brain regions (Fig. 3c). Fig.3 illustrates that the most actively expressed genes are associated with D2 cluster (Fig. 3a). *Ppp1r1b* and *Penk* genes with more than 1000 FPKM units may be considered as the signature, driver genes within D2 cluster. The selected genes in D1 passive cluster also maintain a dorsal striatum-specific expression pattern (Fig. 3b), but their average expression rate was lower than that of D2 cluster genes (Fig. 3a), implying the major role of D2 cluster, and DARPP-32 in particular as the basic maintenance unit of dorsal striatum functioning.

**Figure 3.**
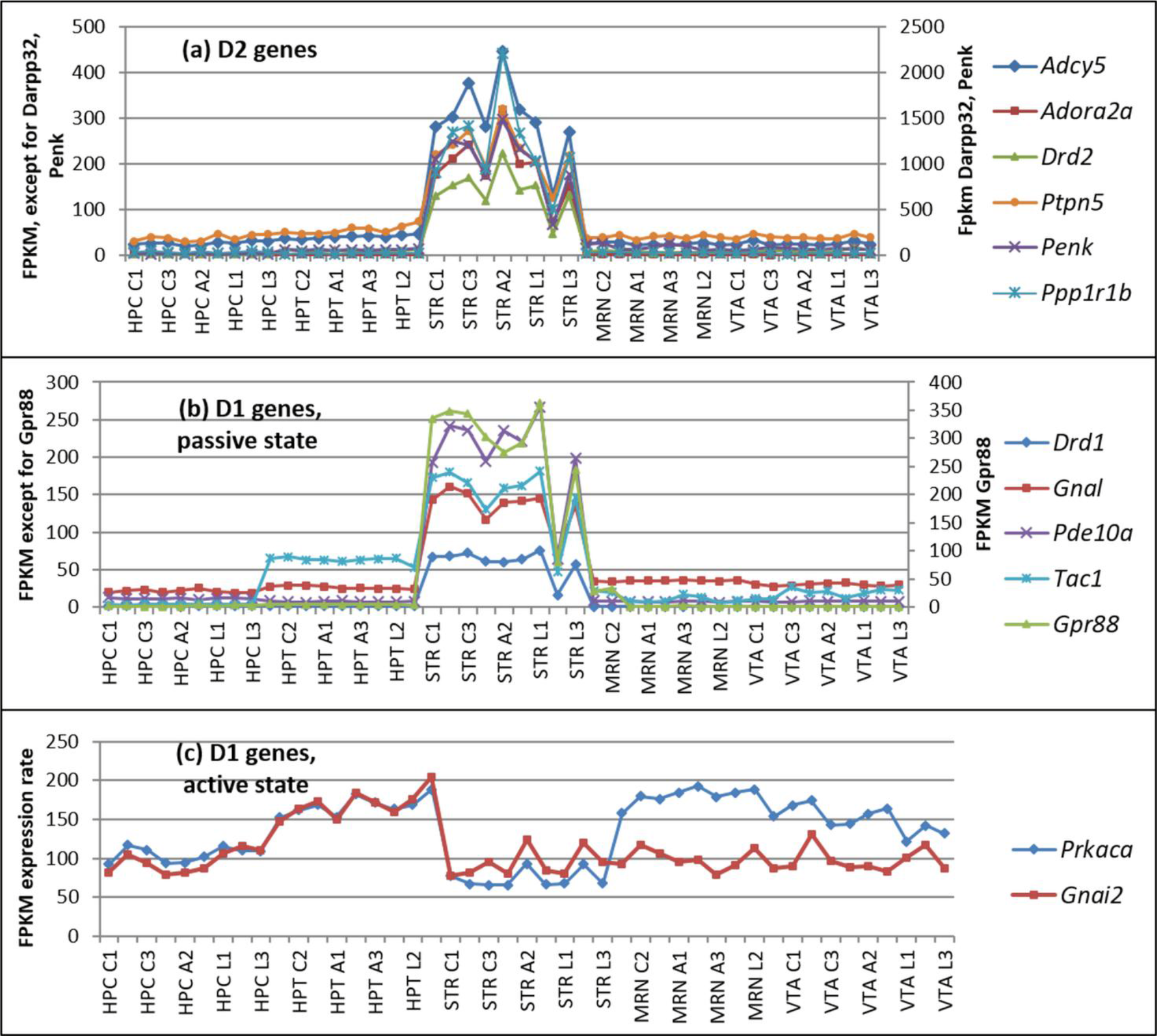
Profiles of D1/D2 striatum specific genes expression (a,b), and non (anti) -specific ones in D1-active pathway (c) across 5 brain regions.C1, C2, C3 – control; A1, A2, A3 – aggressive mice; L1, L2, L3 – losers, defeated mice. HPC-Hippocampus, HPT – hypothalamus, STR-dorsal striatum, MRN – midbrain raphe nuclei, VTA – ventral tegmental area.

Notably, in the above-mentioned two clusters (D2 and D1 passive), the expression rate of selected genes was significantly higher in dorsal striatum compared to other four brain regions. On the contrary, assessment of the signature genes (*Prkaca, Gnai2*) of D1-active cluster demonstrated patently decreased expression rates of *Prkaca* and *Gnai2* genes as compared to other brain regions (Fig. 3c), implying a short expression time span upon firing as one of the reasons.

### Alternative splicing analysis

We retrieved 13 splice isoforms for 12 genes from our set of 33 core genes (Supplemental Table S3). After that we performed Principal Components Analysis (PCA) for the Pearson pairwise correlation matrix. A circular plot for 46 resulting transcripts is shown in Fig.4. Most splicing isoforms, namely for genes *Prkaca, Adora1, Mapk7, Mapk1, Tac1, Drd1, Penk, Ppp3ca* (Fig. 4), exhibit concordant expression patterns in the same clusters or close to them. However, we identified negatively correlated, mutually exclusive splicing isoforms for three pairs of gene transcripts: *Pde10-Pde10_1, Ptpn5-Ptpn5_1, Gnal-Gnal_1* (Fig. 4). Notably, *Pde10a_1, Ptpn5_1* and *Gnal_1* represent minor long isoforms (Supplemental Fig. S1a-c) with negligible expression rates (4-100-fold lower; Supplemental Table S3) as compared to corresponding major transcripts. According to the results shown in Fig.4 one may suggest that the transition from D1 passive to D1 active state is accompanied by the switch between transcription patterns of *Pde10 and Gnal* genes. Also, the transition from D2 state to dopamine depleted state is signified by the switch between transcription patterns of *Ptpn5*.

**Figure 4.**
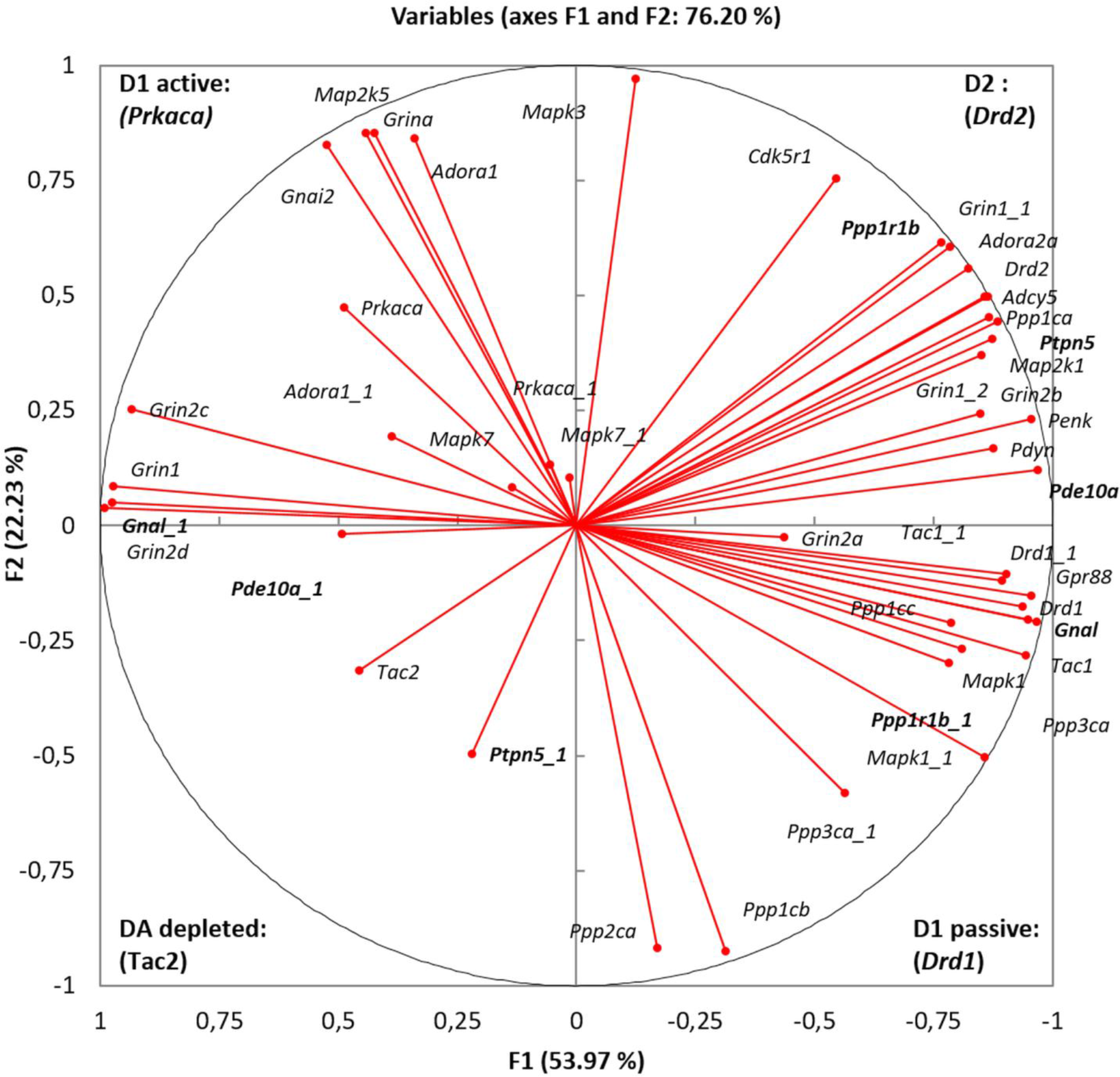
Distribution of splice isoforms across D1/D2 clusters. In bold are splice isoforms that are negatively correlated, or significantly differed (bold italic) in their location. Transcript denotations are listed in Supplemental Table S3.

The *Ppp1r1b* minor splice variant (*t-DARPP* or *Ppp1r1b*_1 in Supplemental Table S3) was found to be specific to D1 MSNs in a passive mode and to exhibit 5-fold reduction in expression level ascompared to that of major D2 cluster-associated transcript (Fig. 4). The truncated isoform of *Ppp1r1b* (*t-DARPP;* NM_001313970; Avanes et al. 2019) (Supplemental Fig. S2a) lacking 37aa at *N*-terminal makes T34 inaccessible to phosphorylation by PKA as well as prevents its overall binding to PP1 while retaining the ability to inhibit PKA upon T75 phosphorylation.

#### Role of CpG islands in alternative transcription

The genes listed above (*Ppp1r1b, Ptpn5, Pde10a* and *Gnal*) are CpG islands associated with promoter genes. Remarkably, upon transition of D1 passive to D1 active state and consequent gene networks rearrangement, some of the alternative long isoforms start being transcribed from its closest distal CpG islands (Supplemental Fig. S1, S2). This alteration is witnessed by negative correlation of the splicing isoforms. This is the case with *Pde10a*, *Gnal* genes (Supplemental Fig. S1), implying that the mechanism of transcription pattern switching is based on the inhibition of proximal promoter CpG island of the major isoform transcript. Two other genes, *Ppp1r1b* and *Ptpn5,* maintain CpG promoter in the major isoform but with non-CpG mediated alternative transcription start site (Supplemental Fig. S1, S2). As the above-mentioned genes are involved in D1 passive/active state dynamics, CpG island inhibition process may support the propensity for rapid reciprocity upon D1 active state abrogation and the restoration of the genes’ default isoforms.

## Discussion

The dynamics of MSN related cAMP-mediated dopamine response, even though intensively studied, needs to be elaborated further (Yapo et al. 2017). After the emergence of the single cell transcriptome protocol, in the study by Gokce et al. 2016 and in a range of subsequent publications dedicated to single cell analysis the taxonomy of D1/D2 neurons and glial cells in dorsal striatum was depicted. Still, the current study showed that the pooled tissue analysis may yield a unique insight based on the consideration of specific genes subsets.

We pursued the task of elucidating the phase portrait of dopamine cAMP-mediated response based on specific genes subset expression profiles to assess the asymptotic steady states of MSNs in the dorsal striatum, taking into account the fact that the dorsal striatum cell content is highly homogeneous. It was postulated that cell identity is determined by specific gene expression signature as it is in single cell analysis. In our work we found four distinct gene clusters, namely: one corresponding to D2 MSNs, another one - to dopamine depleted state, and two clusters - to opposite D1 passive/active states. The analyzed clusters feature the following properties:

1. **D2 cluster.** While it is known that *Ppp1r1b is* expressed both in D1- and D2 MSNs, in the current study the expression rate of *Ppp1r1b* proved to be strongly correlated with those for *Cdk5 (p35)* along with *Drd2, Adora2* and *Penk*, implying that a major fraction of *Ppp1r1b* expression rate in dorsal striatum has to be ascribed to D2 MSNs. Notably, phosphorylation of DARPP-32 and D1R by CDK5 has been reported previously (Jeonget al. 2013. Expression of *Ppp2ca, Ppp3ca* genes encoding the catalytic subunits forspecific serine-/threonine- protein phosphatases PP2A, PP2B was not found in this cluster by the clustering analysis (Fig. 1, 2). Instead, the previously reported tyrosine phosphatase STEP encoded by *Ptpn5* is intensively expressed in the neurons of this type (Gerfen and Surmeier 2011). In D2 MSNs *Adora2A-Drd2-Adcy5* genes constitute a highly interlinked heteromeric transmembrane receptor complex elucidated recently and featuring D2-MSNs (Ferré et al. 2018; Navarro et al. 2018).
2. **D1 cluster (passive phase).***Ppp2ca* (PP2A) and *Ppp3ca*(PP2B) were found to be specifically expressed in D1-neurons, apparently exemplifying dephosphorylation of DARPP-32 upon activation signal abrogation, thereby maintaining homeostasis of phosphorylated DARPP-32 proteins at a certain level mediated by AMPA/NMDA Ca2+ input rate. Also, experimental findings (Massart et al. 2009) that *Golf*, *Gpr88* are the major G-proteins facilitating preferential maintenance of *Drd1* receptors in MSNs have been confirmed. As mentioned earlier (Sin et al.2008) preprotachykinin A (*Tac1) e*xpression is highly correlated with that of D1 receptor. As follows from correlation analysis, ERK2(*Mapk1)* kinase is also involved in maintaining D1 MSN passive phase and was shown to be associated with D1 neurons (Gerfen et al. 2002). Notably, the truncated form of *Ppp1r1b* (*t-DARPP*) was previously reported for several brain structures (Straccia et al. 2016), but it is the first time that *t-DARPP* has beenmappedto MSNs network D1 cluster.
3. **D1 cluster (active phase).**We observed coordinated expression of *Prkaca, Grina* and *Adora1a* genes in active D1 MSNs. Their relation to D1 neurons was reported previously (Svenningssonet al. 2004), along with *Grin1*, *Map2k1*/*Mapk3*(*ERK1*) and *Map2k5*/*Mapk7*(*ERK5*) genes (Grissom et al. 2018), which were also localizedin this cluster in our study.
4. The fourth cluster was represented by single gene, *Tac2* (Fig. 2) which maintains minimal striatal FPKM level of 0.5-1 units observed in the current study. Preprotachykinin B (*Tac2*) is known to be expressed in only 5% of dorsal striatum non-MSN neurons (Zhou et al. 2004). It implies that *Tac2* gene may be chosen as a signature indicator of non-D1/D2 MSN neuron type lacking specific dopamine response, thus indicating the dopamine-depleted state (Fig. 2b).

The expression profiling of the genes involved in D1/D2 MSNs phosphorylation cascades has proved informative and concordant with our current knowledge of gene pathways in these neurons. As an illustration in Fig. 5a, b the proteins ascribed in our study to the genes from D1/D2 clusters are marked with color. This technique and the schematics proposed by (Yapo et al. 2017) can be helpful for gaining the expression-based insight into the specifics of MSNs functioning.

**Fig. 5.**
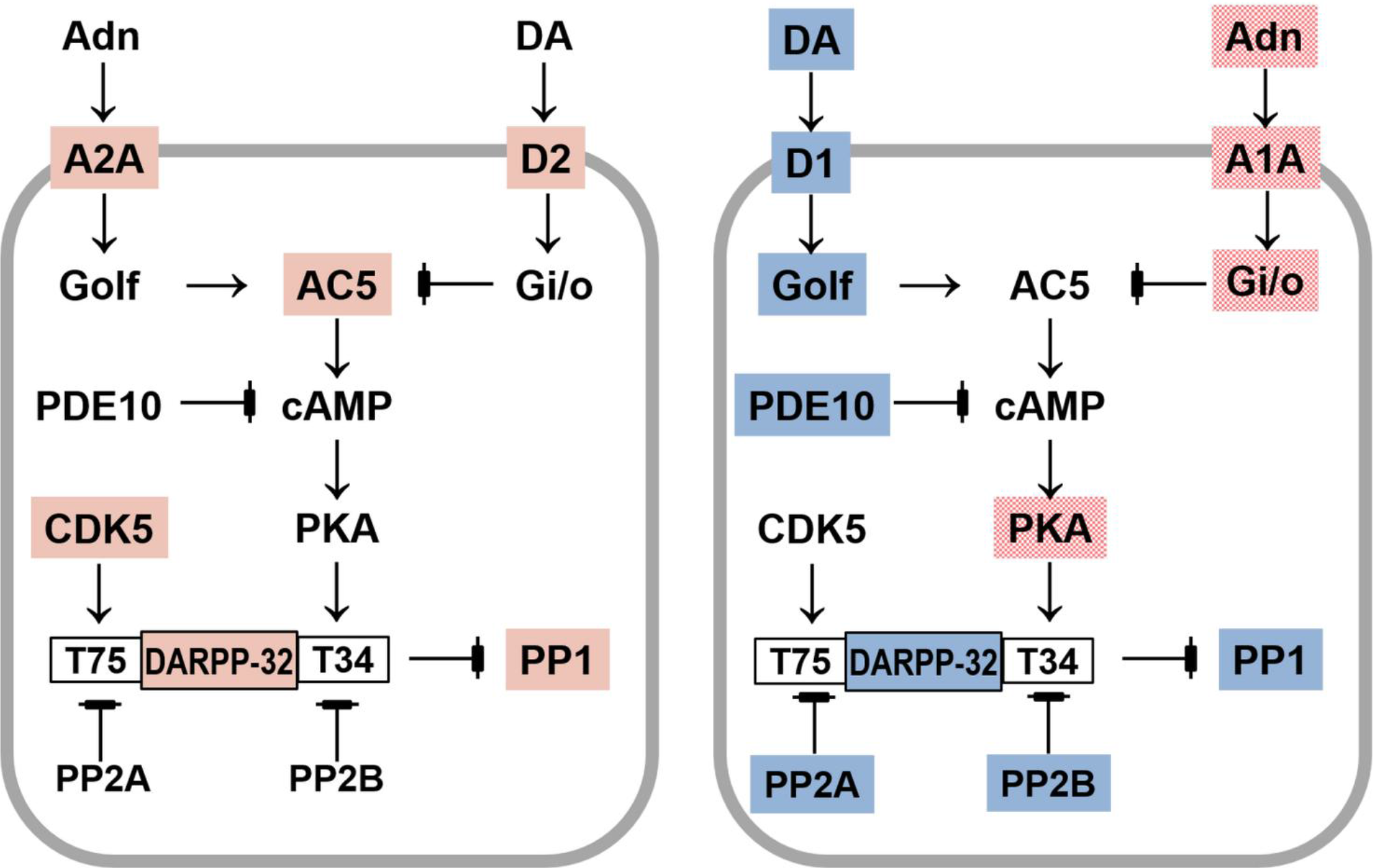
(a). Stable D2-A2A-Golf-Gi/o complex features preferential inhibition of AC5 –> cAMP synthesis upon DA (dopamine) signal, though stimulants can invoke it (Ferré et al. 2018; Navarro et al. 2018). Genes encoding proteins shaded red are coordinately regulated in D2 neurons as observed in our RNA-Seq data (Fig.1). Genes encoding uncolored proteins are absent in the D2 cluster (Fig.1), implying they may be present with minor expression ranges and are not involved in coordinated variation. (b). Oscillating passive-active cascades in D1 neurons. Genes encoding proteins of the same color manifest correlated clusters in our data presented in Fig. 2. Blue (D1 passive state) vs Red (D1 active state) clusters are antagonistic ones according to AHC and PCA analyses (Fig.1; Fig. 2). Genes encoding uncolored proteins are absent in the D1 cluster (Fig.1), implying they may be present with minor expression ranges and are not involved in coordinated variation.

While the variation among the groups of experimental animals yielded no particular group-specific clustering (Fig.2), the observed samples obtained from defeated mice (L2, L3) exhibit a bias toward the dopamine-depleted quadrant (Fig. 2). Aggressive mice (A2- winner) display an increase in both D1 and D2 expression (Fig. 2). Other samples tend to maintain passive D1 phase and moderate D2 MSNs genes expression (Fig. 2). Thus, it can be stated that a behavior model featuring distinct physiological states is a potent tool to enhance variation in expression profiles of observed genes and to ascertain dopamine-depleted area specific states (D1 active, DA depleted). Graphically it is represented in Fig. 2 by blue arrow underlined dopamine-glutamate gradient.

The ‘up’ and ‘down’ states of D1 neurons were reported previously (Wickens and Wilson 1998; Surmeier et al. 2007): the ‘firing’ of D1 neurons upon glutamate input is reported to maintain peak-like induction of voltage increase spanning about 1 sec (Wickens and Wilson 1998) incomparable with the down state span for the major time lapse. Stress model used in the study helped to visualize the D1 active state in detail.

The performed correlation analysis was based on the previous experimental evidence that the synaptic genes network expression is highly coordinated and specific (Trinidad et al. 2008). Our attempt to shed a light on such intricate genes networks was based on the presumption that proteins phosphorylation/ dephosphorylation cascades provide relevant feedback for genes expression rate, which was found to be well-grounded in our study.

As it is known, besides dopamine, there are multiple other neurotransmitters that regulate the excitability of dopaminoceptive neurons: glutamate, GABA, opiates, and adenosine, which are involved in signaling through DARPP-32 in medium spiny neurons (Nishi et al. 2017). In particular, in our previous studies the expression rates of catecholaminergic, glutamatergicand GABAergic receptors in the dorsal striatum were shown to be significantly altered as compared to the control group, in reverse directions for the defeated and aggressive mice (Smagin et al. 2018; Smagin et al. 2019).

DARPP-32 serving as a PKA inhibitor upon Thr75 phosphorylation by CDK5, as reported elsewhere (Bibb et al. 1999; Nishi et al. 2002; Nishi et al. 2017) proved to be specific to D2 neurons. According to our data, the genes with dorsal striatum expression profiles most correlated with that of *Ppp1r1b Cdk5r1* along with *Ppp1ca* (Fig. 2) in D2 cluster. For D1 passive state *Ppp1cc* exhibits a distinct expression pattern (Fig. 2) implying also that *Ppp1r1b* encodes the PP1 subunit.

Notably, phosphorylation of DARPP-32 Thr34 cascade upon PKA activation (Nishi et al. 2017) apparently leads to PP1 phosphatase depletion according to the absence of *Ppp1\** genes encoding the PP1 subunit in D1 active cluster (Fig. 2). Consequently, theinhibition of PP1 by DARPP-32Thr34 phosphorylation in D1 active state upon PKA activation appears to be a fairly rapid and effective process.

Specific combinatorial usage of NMDA receptor subunits (*Grin2a*, *b, c, d*) with content alterations across various clusters (Fig. 2) has been found within the dopamine-dependent cAMP-related genes network.

When analyzing alternatively spliced transcripts, we found a wide range of genes employing alternative promoter usage (Supplemental Fig. S1). Most of them are associated in particular with D1 neurons (*Drd1, Gnal, Ppp1r1b, Pde10a;* Supplemental Table S3; Supplemental Fig. S1, S2), implying a possible mechanism for gene/transcripts switching through reversible blocking of CpG island promoters (Supplemental Table S3; Supplemental Figs. S1, S2). It is worth noting that the minor isoform is usually longer (Supplemental Table S3), implying the inhibition of proximal CpG island, as reported earlier (Sarda et al. 2017). As the switching between D1 active – passive states is of oscillatory nature, CpG islands are apparently not modified by DNA methylation as was the case in (Sarda et al. 2017), nor are alternative histone modifications (Pal et al. 2011), since both isoforms are present in D1 neurons. It is also intriguing that the majority of the isoforms encode alternative functional proteins, which are expressed in D1 active state, but usually at significantly lower rates (Supplemental TableS3).

Notably, D1 MSN excitation kinetics exerts the major impact on the D1/D2 cascades dynamics as follows from D1-split states observations (featuring PKA cascade activation), while D2 neurons manifest the strongest coordinated background expression of the target genes, including postsynaptic *Drd2- Adora2- Adcy5* receptor complex, *Ppp1r1b*, and *Penk* (Fig.2). In particular, *Drd2* expression rate in dorsal striatum is twofold higher than that of *Drd1* (Supplemental Table S1). High *Penk* expression rate observed in D2 neurons (Fig. 3a; Supplemental Table S1) pinpoints D2 MSNs entity as a major neuroendocrine response body in dorsal striatum previously reported in (Dunn et al. 2012; Bali et al. 2015; Belujon and Grace 2015).

In conclusion, we correspond that using RNA-Seq data as the basis for an asymptotic stationary model of gene pathways we were able to elucidate the cAMP-mediated phosphorylation activation pathways. Dopamine response gene pathways specific to D1- and D2-neurons represent the tight clusters of highly correlated genes based on the expression rate profiles. In Fig.5a the signature genes elucidated for the D2 cAMP cascade by cluster analysis are depicted (Fig. 1). Moreover, the clustering was effective in distinguishing the passive and active states of D1-neurons which employ distinct gene non-overlapping pathways instantiation, thereby implementing the oscillatory states of D1 neurons (Fig. 5b).

According to our data, *Ppp1r1b* proved to be one of the most highly expressed protein-encoding genes in dorsal striatum MSNs,. Based on the correlation analysis, the major activities of DARPP-32 in dorsal striatum MSNs are assumed to be related specifically to D2 neurons, including, besides the regulation of PKA activity, the involvement in neuroendocrine stress response. It has to be stressed that the pathways reconstruction attempt succeeded due to the unexpectedly high coordination of the sampled genes expression, as well as the homogeneity/dominance of MSNs content in dorsal striatum. In PCA analysis the overall variation for the first two components reached 86% (Fig. 2), which underscores that our model presentation is exhaustive and consistent. We believe that the stationary phase expression profile provided for dopamine cAMP-mediated response in MSNs will help gaining further insight into the MSNs molecular dynamics.

## Materials and methods

### Animals

Adult C57BL/6 male mice were obtained from Animal Breeding Facility, Branch of Institute of Bioorganic Chemistry of the RAS (ABF BIBCh, RAS) (Pushchino, Moscow region). Animals were housed under standard conditions (12:12 hr light/dark regime starting at 8:00 am, at a constant temperature of 22+/− 2°C, with food in pellets and water available *ad libitum*). Mice were weaned at three weeks of age and housed in groups of 8-10 in standard plastic cages. Experiments were performed with 10-12 week old animals. All procedures were in compliance with the European Communities Council Directive 210/63/EU of September 22, 2010. The protocol for the studies was approved by Scientific Council No 9 of the Institute of Cytology and Genetics SD RAS of March, 24, 2010, N 613 (Novosibirsk, http://spf.bionet.nsc.ru/).

### Generation of alternative social behaviors under agonistic interactions in male mice

Prolonged negative and positive social experience, social defeats and wins, in male mice were induced by daily agonistic interactions (Kudryavtseva 1991; Kudryavtseva et al. 2014). Pairs of weight-matched animals were each placed in a steel cage (14 × 28 × 10 cm) bisected by a perforated transparent partition allowing the animals to see, hear and smell each other, but preventing physical contact. The animals were left undisturbed for two or three days to adapt to new housing conditions and sensory contact before they were exposed to encounters. Every afternoon (14:00-17:00 p.m. local time) the cage lid was replaced by a transparent one, and 5 min later (the period necessary for individuals’ activation), the partition was removed for 10 minutes to encourage agonistic interactions. The superiority of one of the mice was firmly established within two or three encounters with the same opponent. The superior mouse would be attacking, biting and chasing another, who would be displaying only defensive behavior (sideways postures, upright postures, withdrawal, lying on the back or freezing). As a rule, aggressive interactions between males are discontinued by lowering the partition if the sustained attacks have lasted 3 min (in some cases less), thereby preventing the damage of defeated mice. Each defeated mouse (loser) was exposed to the same winner for three days, while afterwards each loser was placed, once a day after the fight, in an unfamiliar cage with an unfamiliar winner behind the partition. Each winning mouse (winners, aggressive mice) remained in its original cage. This procedure was performed once a day for 20 days and yielded an equal number of the winners and losers.

Three groups of animals used were: 1) Controls – mice without a consecutive experience of agonistic interactions; (2) Losers – chronically defeated mice; (3) Winners – chronically aggressive mice. The losers and winners with the most expressed behavioral phenotypes were selected for the transcriptome analysis. The affected mice and the control animals were simultaneously decapitated. The brain regions(hippocampus (HPC), hypothalamus(HPT), dorsal striatum(STR), midbrain raphe nuclei (MRN), ventral tegmental area (VTA) were dissected by one experimenter according to the map presented in the Allen Mouse Brain Atlas [http://mouse.brain-map.org/static/atlas]. All biological samples were placed in RNAlater solution (Life Technologies, USA) and were stored at −70°C until sequencing.

It has been shown in previous experiments (Smagin et al. 2018; Smagin et al. 2019) that the chronically aggressive mice develop motor hyperactivity, enhanced aggressiveness and stereotypic-like behaviors. Chronically defeated mice manifest low motor activity and depression-like behaviors.

### RNA-Seq data collection

The collected brain samples were processed at JSC Genoanalytica (www.genoanalytica.ru, Moscow, Russia). According to the protocol therein, mRNA was extracted using a Dynabeads mRNA Purification Kit (Ambion, Thermo Fisher Scientific, Waltham, MA, USA). cDNA libraries were constructed using the NEBNext mRNA Library PrepReagent Set for Illumina (New England Biolabs, Ipswich, MA USA) following the manufacturer’s protocol and were subjected to Illumina sequencing. More than 20 million reads were obtained for each sample. The resulting “fastq” format files were used to align all reads to the GRCm38.p3 reference genome using the STAR aligner (Dobin and Gingeras 2015) followed by *Cuffnorm* software to assess the expression rate of the transcripts (Trapnell et al. 2012). The identification of alternatively spliced events in the RNA-Seq data was performed with Cuffnorm procedure.

### Statistical analysis

For the transcriptome data, a Principal Component Analysis (PCA) was conducted using the XLStat software package (www.xlstat.com). PCA was based on Pearson product moment correlation matrix calculated on the FPKM value profiles of analyzed genes. We also used a Pearson correlation as a similarity metric for the Agglomerative Hierarchical Clustering (AHC). The agglomeration method comprised an unweighted pair-group average. We avoided using commonly accepted WGCNA method (Langfelder and Horvath 2008) due to its detrimental impact on clustering density because of limited amount of our genes sample (up to 100 transcripts).

## Acknowledgments

Supported by Russian Science Foundation (grant N 19-15-00026)

