## Supplementary Figures for "Dopamine response gene pathways in dorsal striatum MSNs from a gene expression viewpoint: cAMP-mediated gene networks"

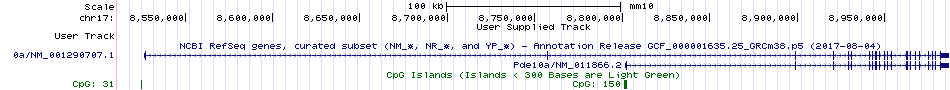

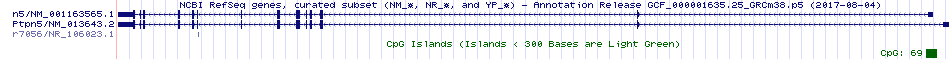

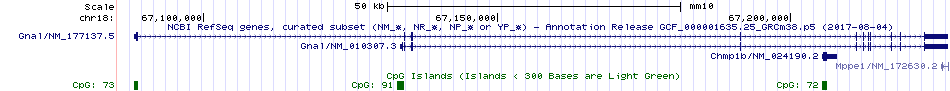

b

c

a

**Supplemental Figure S1.** Mutually exclusive (negatively correlated) splice variants for *Pde10a (a), Gnal (b), and Ptpn5 (c)* schema of the alternative CpG promoters. Corresponding CpG islands are encircled.

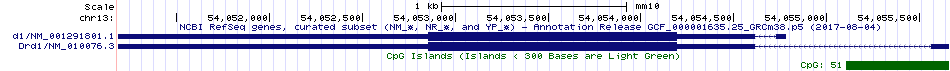

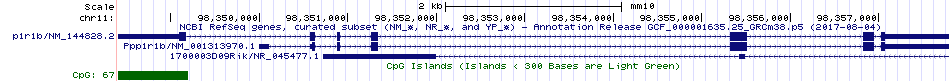

a

b

**Supplemental Figure S2.** Structure of alternative isoforms of two genes *Ppp1r1b* (DARPP-32) (a), *Drd1* (b) based on their alternative (non-CpG) promoters.
